# A vascularized 3D model of the human pancreatic islet for *ex vivo* study of immune cell-islet interaction

**DOI:** 10.1101/2021.12.21.473744

**Authors:** R. Hugh F. Bender, Benjamen T. O’Donnell, Bhupinder Shergill, Brittany Q. Pham, Damie J. Juat, Michaela S. Hatch, Venktesh S. Shirure, Matthew Wortham, Kim-Vy Nguyen-Ngoc, Yesl Jun, Roberto Gaetani, Karen L. Christman, Luc Teyton, Steven C. George, Maike Sander, Christopher C.W. Hughes

## Abstract

Insulin is an essential regulator of blood glucose homeostasis that is produced exclusively by β cells within the pancreatic islets of healthy individuals. In those affected by diabetes, immune inflammation, damage, and destruction of islet β cells leads to insulin deficiency and hyperglycemia. Current efforts to understand the mechanisms underlying β cell damage in diabetes rely on *in vitro*-cultured cadaveric islets. However, isolation of these islets involves removal of crucial matrix and vasculature that supports islets in the intact pancreas. Unsurprisingly, these islets demonstrate reduced functionality over time in standard culture conditions, thereby limiting their value for understanding native islet biology. Leveraging a novel, vascularized micro-organ (VMO) approach, we have recapitulated elements of the native pancreas by incorporating isolated human islets within a three-dimensional matrix nourished by living, perfusable blood vessels. Importantly, these islets show long-term viability and maintain robust glucose-stimulated insulin responses. Furthermore, vessel-mediated delivery of immune cells to these tissues provides a model to assess islet-immune cell interactions and subsequent islet killing — key steps in type 1 diabetes pathogenesis. Together, these results establish the islet-VMO as a novel, *ex vivo* platform for studying human islet biology in both health and disease.

## INTRODUCTION

Diabetes affects over 34 million individuals in the U.S. and is broadly grouped into type 1 (T1D) typically showing juvenile onset, and type 2 (T2D), which is most common in adults. Both types of diabetes are characterized by hyperglycemia caused by either relative or absolute insulin deficiency (1). Although the exact mechanisms of diabetes pathogenesis remain unclear, inflammation and/or autoimmune reactivity against insulin-producing pancreatic β cells are key drivers in both forms of diabetes (2; 3). During T1D pathogenesis, β cells are destroyed by an autoimmune process, leading to insulin deficiency and subsequent hyperglycemia (4). In contrast, T2D is characterized by insulin resistance in peripheral tissues and relative insulin deficiency caused by chronic damage to β cells (5), which is thought to result, in part, from cytokine-mediated activation of both tissue-resident and circulating macrophages (6; 7). To date, most studies of diabetes pathogenesis have relied on rodent models, yet these models do not account for important differences in islet structure, function, and immunology between human and rodent (8; 9). Thus, there is increasing emphasis on developing better *in vitro* tools that utilize human islets to study diabetes.

During development, endocrine precursor cells co-develop with endothelial cells (EC) such that blood vessels surround and penetrate mature islets in the adult pancreas, allowing for rapid sensing of blood glucose levels and subsequent release of relevant hormones (10–13). β cells exhibit apical-basal polarity relative to these blood vessels (14; 15), and the surrounding laminin-rich extracellular matrix (ECM) supports integrin signaling that drives β cell proliferation as well as insulin expression and release (16; 17).

Standard islet isolation procedures physically disrupt and remove both blood vessels and ECM, leading to reduced glucose responsiveness and cell death within only a few days (18; 19), underscoring the need for new approaches that better maintain isolated human islets. Previously, we developed a microfluidic vascularized micro-organ (VMO) platform (20–22) in which capillary-like vessels deliver nutrients directly to, and remove waste from, the surrounding tissue, just as in the body (23; 24). Here, we present a VMO platform that incorporates human cadaveric islets within a 3D vascularized tissue — the islet-VMO. The platform mimics the native islet niche and β cells receive glucose via the surrounding vasculature, which stimulates insulin secretion comparable to native islets *in vivo*. Moreover, as a proof-of-principle, activated immune cells introduced via the vasculature extravasate and migrate into the islets, mimicking the early stages of T1D disease progression. Collectively, these findings demonstrate the utility of the islet-VMO as a novel biomimetic model for understanding islet biology and diabetes pathology.

## RESEARCH DESIGN & METHODS

### Platform Fabrication

The islet-VMO platform was fabricated using standard PDMS photolithography techniques as previously described (24), except that later devices used 200μm outer channels rather than 100μm channels.

### Islet Isolation

Non-diabetic human cadaveric islets were provided by the NIDDK Integrated Islet Distribution Program (IIDP) or Prodo Laboratories Inc. (Aliso Viejo CA). Upon arrival, islets were washed in islet medium (Table S1) then stained with dithizone. Dithizone^+^ islets <200μm in diameter were hand-selected under a dissection microscope then allowed to recover overnight in islet medium.

### Islet-VMO Loading

Islets were co-loaded (20-25 islets/μL hydrogel) with 7×10^6^ cells/mL of endothelial colony-forming endothelial cells (ECFC-ECs, “EC”) and stromal cells (NHLFs, Lonza Bioscience, Rockville MD) in 8mg/mL fibrin hydrogel clotted with 50U/mL thrombin. Islets were maintained within the islet-VMO for up to two weeks by gravity-driven flow of EGM-2 medium (Lonza Bioscience) through the vasculature (22; 24; 25).

### Vascular Network Quantification

Vessel networks were perfused with 50µg/mL 70kDa FITC or rhodamine-dextran and imaged on a Nikon Ti-E Eclipse inverted fluorescent microscope. Morphological measurements of perfused vessels were collected using FIJI imaging software by tracing the mid-point of all vessels through the cell chamber (Fig. S1). Branch points were quantitated at the intersection of these mid-point lines. Lastly, vessel diameter was determined by measuring across each traced vessel at ∼50µm intervals.

### Immunofluorescence Staining

Islets were fixed (4% paraformaldehyde), dehydrated, embedded in OCT mounting medium, and 10µm sections were collected. Sections were incubated with primary antibodies (Table S2) overnight at 4°C, then with species-appropriate secondary antibodies, followed by DAPI counterstaining. Islet-VMO platforms were perfused with 4% paraformaldehyde (30 minutes, 25°C), then flushed with DPBS. The bottom silicon membrane was removed prior to antibody staining. Confocal image stacks of sectioned or device-embedded islets were acquired on a Leica TCS SP8 confocal microscope.

### Static glucose-stimulated insulin secretion (GSIS)

GSIS assays were performed to confirm islet function. Recovered islets were starved in low-glucose Krebs-Ringers Buffer (KRB, Table S1) for one hour, then incubated in five groups of ten islets per well in either standard (2.8mM glucose) KRB or high-glucose KRB (16.7mM glucose) for 1 hour. Supernatant was collected and frozen for subsequent insulin assay. Islets were also lysed in acid ethanol (4°C) and collected for total insulin. Insulin was measured by ultra-sensitive ELISA (Mercodia, Uppsala Sweden).

### Islet-VMO GSIS

Devices were perfused with M199 medium containing 5.5mM glucose (“Standard M199(+),” Table S1) for one hour. Islets were then stimulated for one hour by adding M199(+) containing 16.7mM glucose (“High-Glucose M199(+)”) to reservoir V_A_ (Fig. 1A), followed by Standard M199(+) for two hours to allow islets to return to baseline insulin secretion levels. M199(+) supplemented with 30mM KCl was then perfused for one hour to assay for releasable insulin. Accumulated device flow-through was collected continuously at 10-minute intervals beginning thirty minutes prior to glucose stimulation. Glucose concentration in the flow-through was assessed using a Contour glucometer and insulin was measured by standard insulin ELISA (Mercodia).

**Figure 1.**
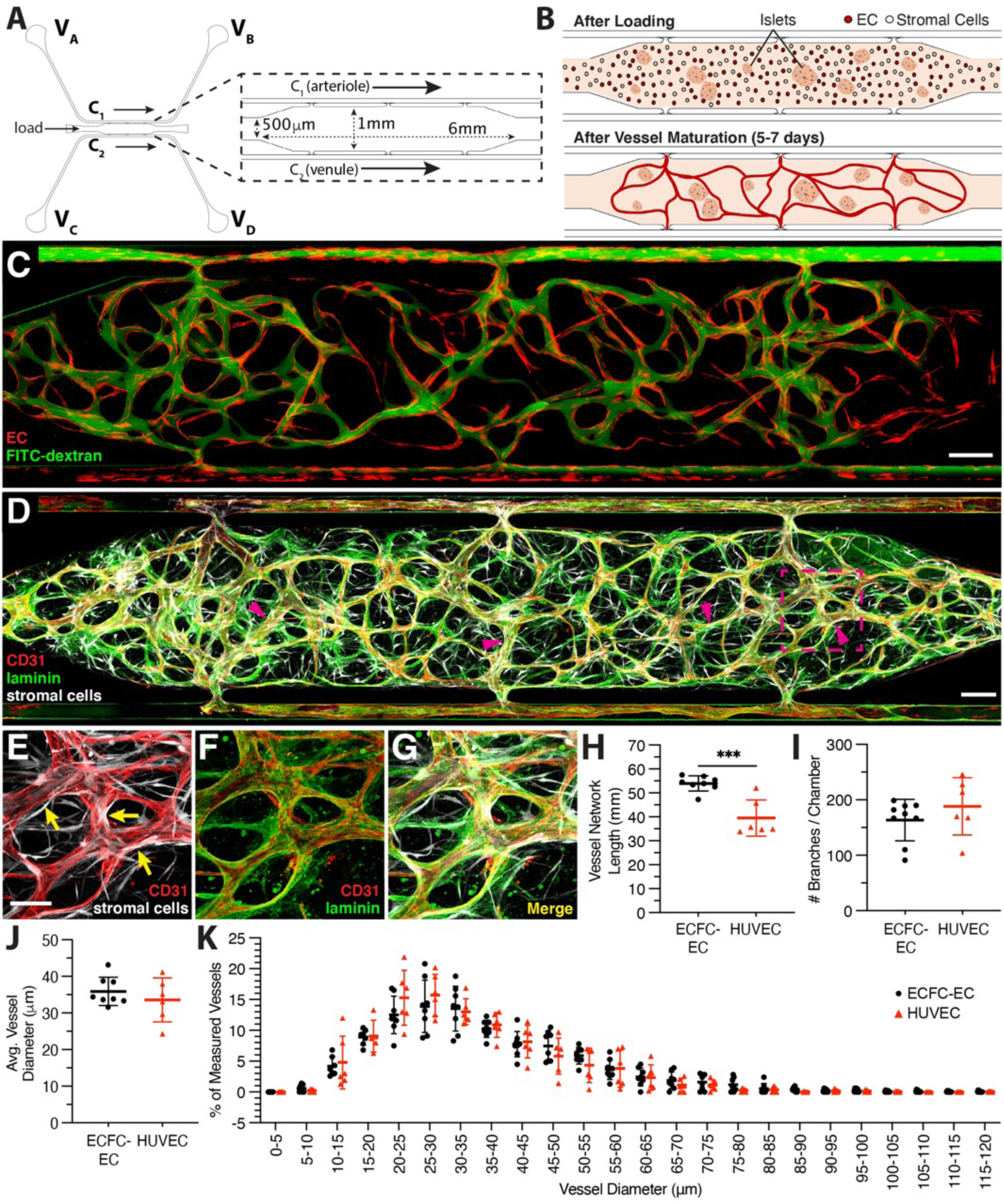
Generating blood vessels in an islet-specialized Vascularized Micro-Organ (islet-VMO). (A) A microfluidic design for generating vascularized islet tissues comprises a central, enlarged cell chamber (inset) flanked by two channels, C_1_ (acting as an arteriole) and C_2_ (venule). Reservoirs at the end of each channel (V_A_-V_D_) are filled with varying heights of medium such that medium is driven by hydrostatic pressure in a net direction from V_A_, through the tissue chamber, to V_D_. (B) To generate vascularized tissues, islets, EC, and stromal cells are loaded together in a fibrin hydrogel through the loading tunnel (labeled in A). Interstitial fluid flow is driven across the cell chamber to activate vessel formation in the central chamber, with development of intact vessel networks complete by 5-7 days post-loading, at which time convective flow through the vessels becomes dominant. (C) Upon maturation, blood vessels (fluorophore-transduced ECs, red) carry 70kDa FITC-dextran (green) with minimal leak. (D, E, G) Mature vessels are wrapped by stromal (pericyte) cells (magenta arrows; transduced stromal cells, white). (D, F, G) These stromal cells help remodel the hydrogel to generate basement membrane (a-laminin, green) around the CD31^+^ endothelium (red). Scale bar, 200mm, inset, 100mm. (H-K) Vessel morphology was compared using either endothelial colony-forming cell-derived EC (ECFC-EC) or human umbilical vein endothelial cells (HUVEC), as measured by vessel length (H), branching (I), diameter (J), and hierarchical diameter distribution (K). All error bars represent standard deviation, significance determined by Student’s t-test (significance ***p<0.001).

### GSIS Mathematical Modeling

Mathematical modeling was performed in COMSOL Multiphysics (COMSOL, Inc., Burlington MA). 2D models of the islet-VMO were created in AutoCAD by tracing experimental images of FITC-dextran perfused vessels. Medium flow was modeled using free and porous media flow and laminar flow physics, and the momentum transport modules were coupled with transport of diluted species physics as detailed previously (26) to simulate transport of oxygen, glucose, and insulin. The islets consumed oxygen and glucose, and secreted insulin with previously reported kinetic functions (27). The PDMS was permeable to oxygen and media entering the device was in equilibrium with the atmospheric oxygen (28). Glucose was introduced through the inlet and the insulin and glucose output was measured by using time integral of outlet amount over 10-minute intervals. The constants used in the model are given in Table S3.

### Generating Pseudo-islets

Unsized islets were recovered overnight in islet medium and then digested with Accutase for 12 minutes at 37°C. After gentle trituration the cells were recovered by centrifugation. Islet cells (600,000/well) were seeded into Aggrewell 400 24-well plates (StemCell Technologies, Vancouver Canada), either alone or mixed with EC and stromal cells at a 5:1:1 ratio. Islet function was measured by static GSIS at 2 days post-reaggregation.

### Immune Modeling

Single-donor PBMCs (200,000/well) were cultured for five days in round-bottom, ultra-low attachment 96-well plates (Corning, Corning NY) along with IFN-*γ*-activated islets (10/well) and PBMC activation cocktail (Table S1), in the presence or absence of MHC class I and class II blocking antibodies, 10μg/mL each (Table S2). Resultant cell populations were tested for cytotoxic activity by incubation with donor-matched islets (10 islets/well) for two days. Supernatant was then collected for cytokine analysis by ELISA. Whole islets were also incubated with Hoechst 33342 and live/dead stained.

For perfusion through the islet-VMO, PBMCs were stained with 1µg/mL Cell Tracker Green (CMFDA) dye (Thermo Fisher Scientific) then added to the input well at 5×10^5^ cells/mL. PBMCs either in the vessels (adherent) or in the extravascular space (extravasated) were quantified. Extravasated PBMC within the outline of an islet (invasive), or within 100μm of the islet (islet-adjacent) were counted. These regions of interest were then copied and pasted to areas devoid of islets and PBMC in these areas were quantified to give a value for background PBMC extravasation.

### Statistical Analysis

The Grubbs outlier test was used to determine statistical outliers within sample groups. Statistical significance (*p<0.05, **p<0.01, ***p<0.001, ****p<0.0001) was determined using either an unpaired, two-tailed Student’s t-test or ANOVA, as appropriate, using GraphPad Prism 9 software (GraphPad Software, La Jolla CA).

## RESULTS

### Blood vessel formation in the islet-VMO platform

To generate an *ex vivo* endocrine pancreas model, we leveraged our VMO technology to create a 3D tissue with living, perfusable blood vessels that supply nutrients to the embedded islets. The platform consists of two microfluidic channels, C_1_ and C_2_ (Fig. 1A), flanking a central tissue chamber. To generate blood vessels, hydrogel containing EC and stromal cells is loaded from the side tunnels into the central cell chamber. To form the islet-VMO, islets are loaded together with the blood vessel-forming cells (Fig. 1B). Once vessels form ∼5-7 days post-loading, a blood-substitute medium is driven by a hydrostatic head into C_1_ (the functional arteriole), through the microvascular network, and out through the functional venule, C_2_ (Fig 1B).

When perfused with 70kDa FITC-dextran, the vessel networks demonstrated minimal leak, as indicated by a high fluorescent signal within the vessels and absence of fluorescent signal in the extravascular space (Fig. 1C). *In vivo*, a feature of stable microvessels is the presence of pericytes that wrap around the vessels and promote a mature phenotype. As we have noted before (24; 29) a proportion of the stromal cells we add can differentiate into pericytes, and we see that also in the islet-VMO (Fig. 1D-E). Additionally, the presence of a laminin-rich basement membrane—a key hallmark of microvessels—is found proximal to the abluminal vessel membrane of CD31^+^ blood vessels (Fig 1D-G).

Importantly, ECs from different sources formed similar networks in the islet-VMO. With either ECFC-EC or human umbilical vein EC (HUVEC), networks formed that were perfusable with 70kDa FITC-dextran and showed comparable morphology (Figs. 1H-J, S1) and a similar distribution of vessel diameters (Fig. 1K). While we chose to use ECFC-EC for our studies, both EC sources are suitable in the islet-VMO.

### The islet-VMO platform supports islet vascularization and preserves islet cytoarchitecture

To match the dimensions of our microfluidic device, we size-selected islets <200µm in diameter. While islets of this size constitute the lower 50^th^ percentile of islets in an average donor, evidence indicates that smaller islets (<125µm) contain more β cells, have a higher insulin content, and are more glucose responsive than larger islets (>150µm) from the same donor (30). Islets distributed evenly throughout the tissue chamber (Fig. 2A) and vessel formation was uniform (Fig. 2B). On average, we achieved 20 islets loaded per chamber (Fig. 2C), with an average diameter of 88µm per islet (Fig. 2D).

**Figure 2.**
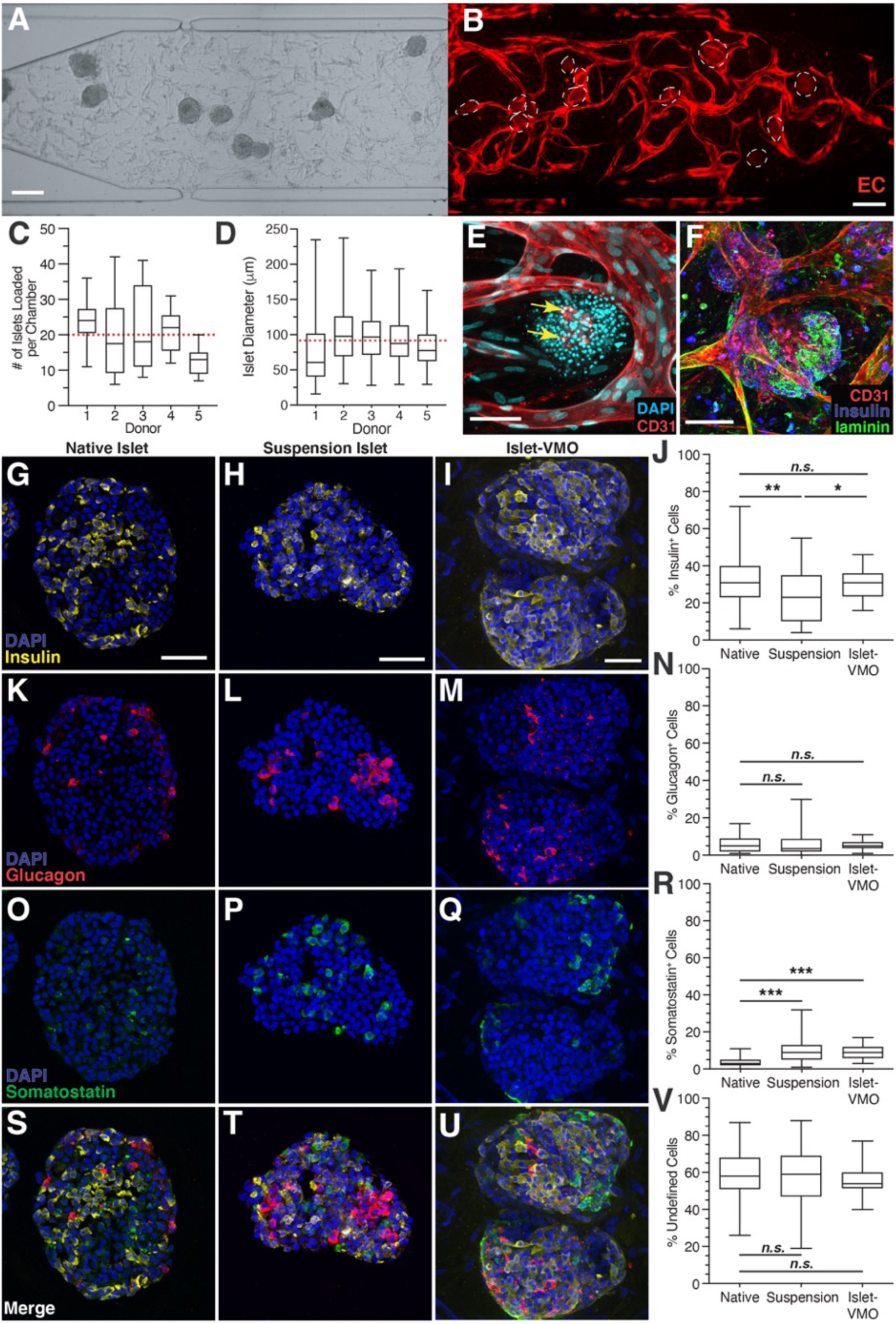
The islet-VMO platform supports islet-proximal vascularization and preserves islet cytoarchitecture. (A) Islets were loaded together with ECs and stromal cells such that an even distribution of all populations is observed after loading. (B) Imaging of the same platform after one week of maturation shows an intact vessel network (red, transduced ECs) around the embedded islets (dashed outline). Scale bar, 200µm. (C) The number of islets was quantified per chamber across five donors, yielding an average of 20 islets loaded per chamber (red dashed line). (D) Quantification of islet diameter across the same islet set shows an average diameter of 87.9µm (red dashed line) (n>110 islets). Box and whisker plots represent median, 25^th^, and 75^th^ percentiles (box) and min and max values (whiskers) for each data set. (E) Immunofluorescent staining for CD31-expressing endothelium (red) shows vessel formation immediately proximal to islets (DAPI, cyan). (F) Immunofluorescent staining of laminin (green) shows the presence of basement membrane surrounding islets (insulin, blue) and CD31+ endothelium (red). Scale bar, 75mm. Immunofluorescent staining and quantification of insulin^+^ b cells (G-J), glucagon^+^ a cells (K-N), and somatostatin^+^ d cells (O-R) was compared between cryosectioned native islets, cryosectioned islets maintained for one week in suspension culture, and *in situ* imaged islet-VMO-embedded islets (n>45 islets across 8 donors). All images were acquired on a confocal microscope and represent maximal projections of the combined image stack. Scale bars, 50mm. (S-U) Merged images of all staining and (V) quantitation of the unstained (non-endocrine) cells in each islet population.

Lumenized vessels formed immediately proximal to islets within 5-6 days of loading, but did not penetrate them (Fig. 2E). The rapid lumenization of ECs next to islets likely results from islet-derived VEGF-A, which is known to be responsible for vascularization both during development and upon islet transplantation (31). Although endogenous CD31^+^ EC fragments are visible in some islets, these fragments do not connect with the surrounding vessels. The islets are, however, surrounded by a laminin-rich basement membrane, just as in the human pancreas (32), despite the absence of laminin in the initial hydrogel (Fig. 2F). Together, these data indicate that the islets promote nearby vessel formation and, in combination with the surrounding stromal cells, remodel the hydrogel to generate laminin-rich basement membrane that recapitulates the native islet environment.

To determine whether the cytoarchitecture of islet-VMO islets is maintained, we compared the proportion of insulin-, glucagon-, and somatostatin-expressing cells in native (freshly-received) islets, islets cultured for one week in suspension culture (“suspension islets”), and islets cultured for one week in the islet-VMO. Suspension islets have fewer insulin^+^ cells compared to native islets, whereas insulin^+^ cells are maintained in the islet-VMO (Fig. 2G-J). Glucagon^+^ cell proportions are similar across all three islet populations (Fig. 2K-N), but the number of somatostatin^+^ cells is increased in both the suspension and islet-VMO islets, relative to native islets (Fig. 2O-R). In all islet populations, the level of undefined (non-endocrine) islet cells remains the same (Fig. 2S-V). Together, these data indicate that the islet-VMO maintains native islet cytoarchitecture and represents an improvement over standard islet suspension culture.

### Glucose responsiveness of islet-VMO islets reflects in vivo islet responses

To determine whether islet-VMO islets respond to glucose stimulation, we performed a GSIS assay by adding glucose-supplemented medium to reservoir V_A_ and collecting effluent from reservoir V_D_ (Fig. 3A). Reservoirs V_B_ and V_C_ were blocked to ensure all glucose flowed through the tissue chamber and that secreted insulin was not diluted by medium cross-flow in the venule channel. In response to perfusion with high glucose (16.7mM) medium we observed a gradual increase in insulin output (Fig. 3B) that returned to baseline once flow of non-stimulatory (5.5mM glucose) medium was restored. As expected, the positive control of KCl stimulation triggered a large increase in insulin release. Interestingly, we noted significant inter-donor variation of insulin secretion profiles. Responses can be slightly delayed (Fig. 3C), more rapid (Fig 3D), peak later (Fig. 3E), or rise from a higher unstimulated baseline (Fig. 3F). Similar donor variation has been documented elsewhere, suggesting that donor islets fall into distinct functional subgroups (33).

**Figure 3.**
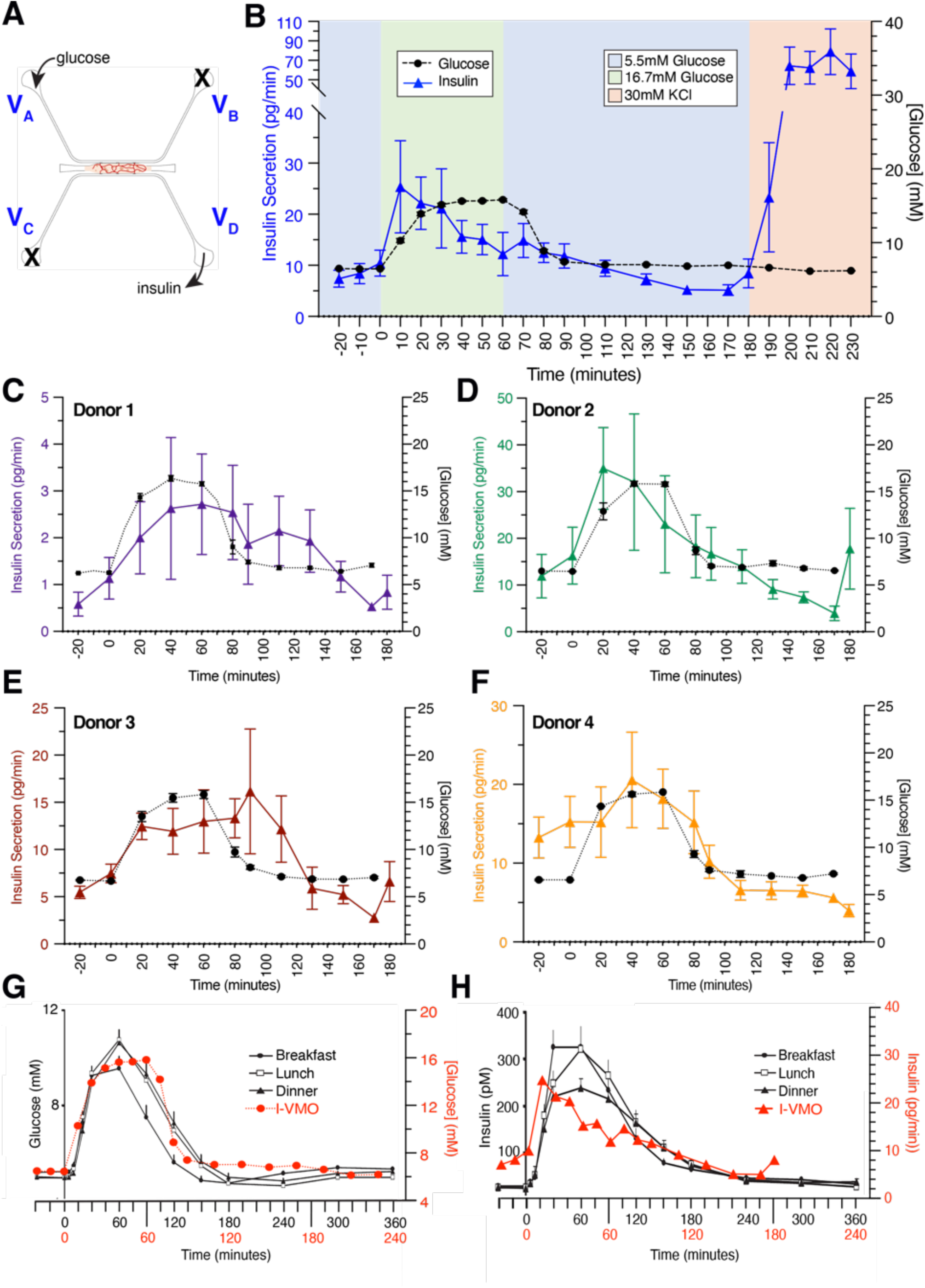
Glucose responsiveness of islet-VMO islets reflects *in vivo* glucose responsiveness. (A) To test islet response to glucose stimulation, medium supplemented with either basal (5.5mM) or high (16.7mM) glucose was introduced through reservoir V_A_ into the vascular network. Insulin and glucose was measured from effluent collected at reservoir V_D_. Reservoirs V_B_ and V_C_ were blocked so that all glucose flowed through the vessel network and secreted insulin was not diluted by cross-flow from V_C_. (B) Perfused glucose (black dashed line) and secreted insulin (blue line) were measured at 10-minute intervals from collected effluent and plotted over time (n=19 Islet-VMOs containing islets from 6 different donors). Error bars represent SEM. (C-F) Measured glucose (black dashed line) and insulin (colored lines) from individual donors show similar secretion patterns despite different magnitudes of insulin secretion (note different y-axis scale) (n≥3 Islet-VMOs per donor). (G) The gradual increase in glucose concentration in the islet-VMO mirrors the dynamics seen *in vivo*. (H) This also holds true for insulin release (islet-VMO data from panel B, human data adapted with permission from (35)).

Importantly, the dynamic insulin secretion response observed in the islet-VMO closely mirrors what is seen in healthy human subjects, whereas this is not the case for the typical profile observed in standard perifusion assays (first phase-second phase (34)). Specifically, we compared aggregated glucose and insulin traces from multiple islet-VMOs to those measured from healthy patients following a meal (Fig. 3G-H, (35)). Overlays show similar dynamics for both glucose and insulin, although the insulin spike is more rapid, less sustained, and occurs over a slightly shorter time frame in the islet-VMO. These differences likely reflect the smaller scale of the platform and the reduced distances traveled by both glucose and insulin relative to distances in the body. Together, these data demonstrate that islets within the islet-VMO respond to glucose stimulation, recapitulate islet donor-to-donor variation, and mimic the coupling of glucose and insulin levels observed in the human body.

### Mathematical Modeling of islet function in the islet-VMO

To better understand the dynamics of glucose and insulin perfusion through the islet-VMO, we employed finite element modeling of actual islet-VMO networks using COMSOL. To model networks, images of FITC-dextran perfused vessels (Fig. 4A) were traced and converted to a two-dimensional model with accurately sized islets positioned at the same locations as in the islet-VMO (Fig. 4B). Using the same hydrostatic pressure heads, glucose concentrations, and time intervals as those tested in the islet- VMO, the mathematical modeling allows estimation of spatial profiles of oxygen consumption, glucose diffusion, and insulin within the islet-VMO (Fig. 4C-E). As expected, we see high levels of oxygen consumption near islets (Fig. 4C), consistent with their known high metabolic activity. Higher glucose concentrations are found closest to perfused vessels, as expected, whereas islets near the periphery receive comparatively less glucose due to their distance from the glucose source – the perfused vessels – and the glucose diffusion rate through the ECM (Fig. 4E). At least some insulin secretion is observed from all islets, however, we note that secreted insulin pools in the vicinity of islets not bounded by a perfused vessel. To validate the mathematical model, we measured actual glucose and insulin concentrations sampled from port V_D_ (from one of the Donor 2 devices – aggregated data shown in Fig. 3D) and compared these to values determined by mathematical modeling of the same vascular network supplemented with islets of the same size and placement (Fig. 4F-G). We note that the sharp insulin secretion spike in the experimental data mirrored the response of islets from this donor set (Donor 2), which responded more rapidly than other sets (see Fig. 3D). Together, these data show that the simulation matches well with the experimental data.

**Figure 4.**
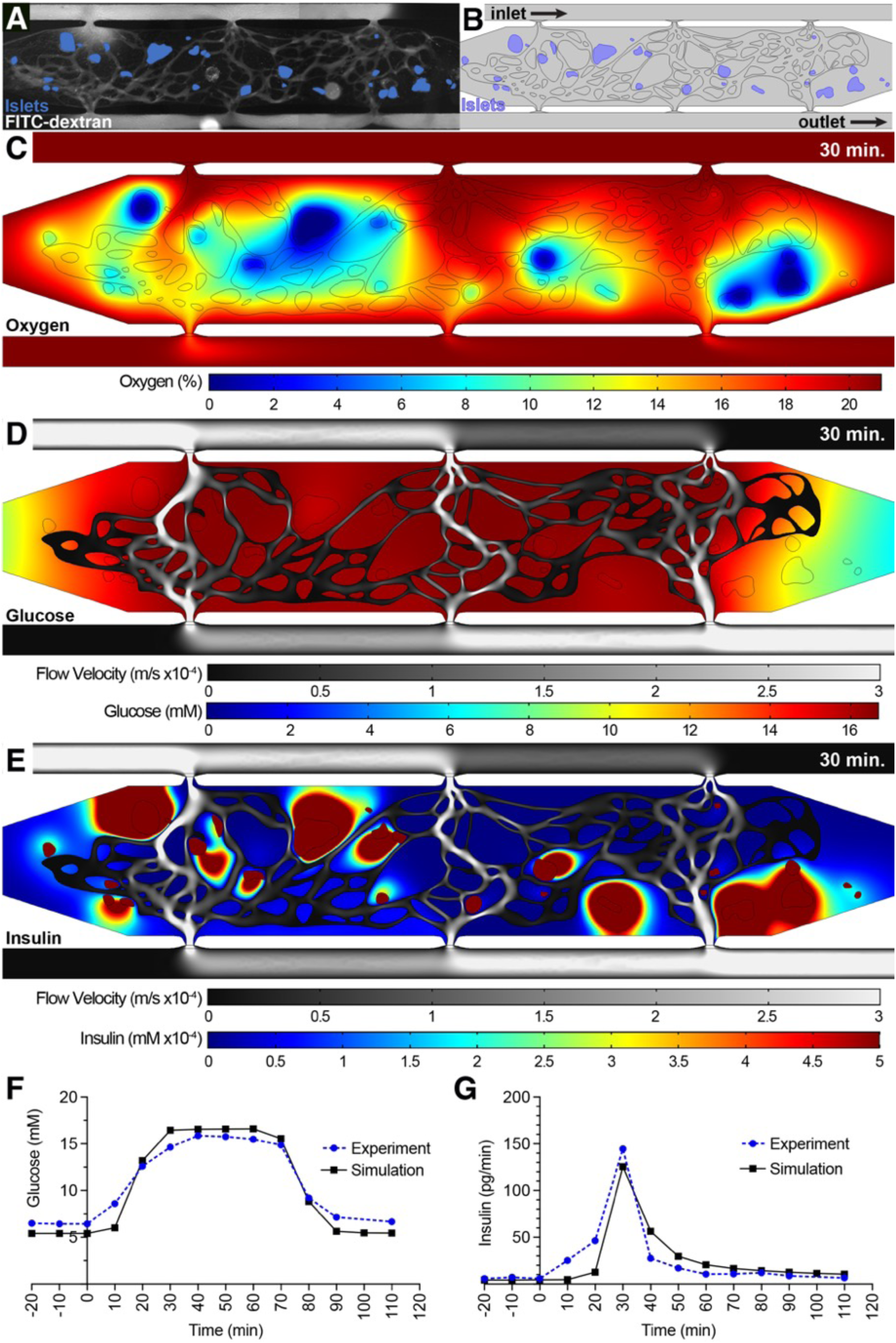
COMSOL modeling of islet function within the islet-VMO platform. (A) Islet-VMO platforms were perfused with FITC-dextran (gray) such that both the perfused vasculature and position of islets within the islet-VMO could be mapped. (B) For COMSOL modeling, vessels were converted to a skeletonized vessel network (black outline) and islets localized to their mapped locations (blue). Glucose perfusion was modeled entering from the inlet at top left with glucose and insulin measurements modeled at the outlet bottom right (equivalent to V_A_ and V_D_ in Fig. 3). Snapshots were acquired from the model at 30 minutes after high glucose perfusion showing (C) islet oxygen consumption (%, color scale), (D) medium velocity (m/s, gray scale) and glucose diffusion (mM, color scale), and (E) medium velocity (m/s, gray scale) and insulin secretion (mM, color scale). (F) Glucose perfusion and (G) insulin secretion traced over time in the COMSOL simulation model (black line) was compared to experimental values from the same islet-VMO (blue line).

As noted above, not all islets secrete insulin at the same rate. To understand how individual islets contribute to overall insulin output, we mathematically shut down insulin secretion from all but one islet, and repeated this to examine islets individually. Analyzed islets were selected based on: (a) their location within the chamber (either in the center [#2-5] or at the periphery [#1, #6]); and, (b) their proximity to perfused vessels (either close [#1-4] or distant [#5-6]) (Fig. 5A). Consistent with the hypothesis that perfused vessels allow for efficient transportation of islet-secreted insulin, islets closest to vessels (#2-4) demonstrate the highest release of insulin into the vessels as measured at the outlet V_D_ (Fig. 5B). However, islets at the periphery where vessels have lower perfusion (#1) contribute minimally to overall insulin output. In addition, islets distant from perfused vessels (#5, #6) also demonstrate both delayed and diminished insulin secretion, independent of their location within the chamber. To further demonstrate the importance of perfusable vessels in proximity to islets for improved insulin clearance, we performed additional modeling on islet #3, in which a new vessel was added to the top side of the islet (Fig. S2A-D). This dramatically reduces the accumulation of insulin around the islet and increases insulin levels measured at the outlet (Fig. S2E-F). Together, these simulation data demonstrate the importance of the vascular network for appropriate coupling of glucose levels and insulin output.

**Figure 5.**
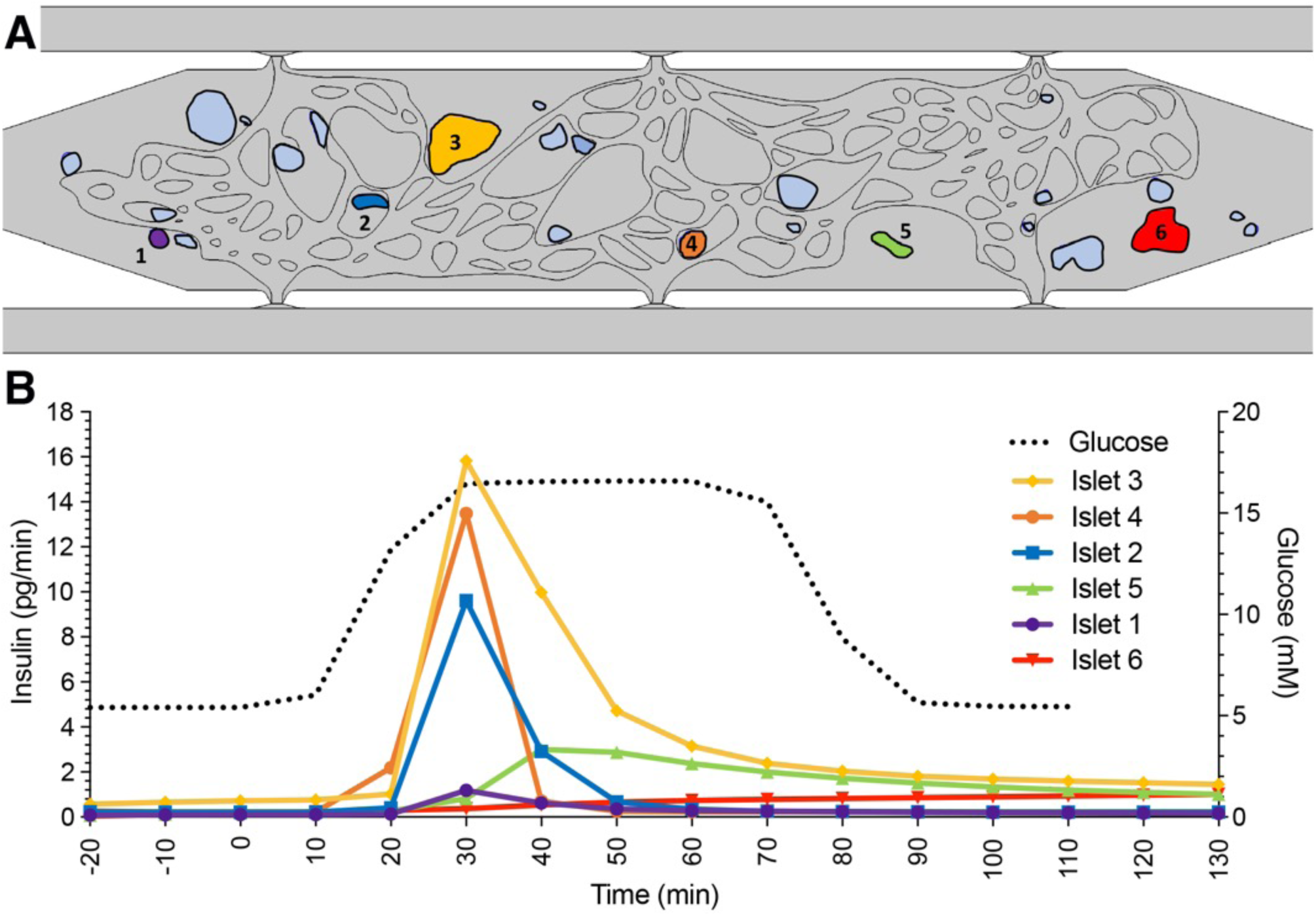
Individual islet function modeled within the islet-VMO. (A) Using the COMSOL model described in Fig. 4, six islets (annotated and colored) were selected for measuring individual islet function. (B) Insulin secretion profile as measured at the outlet V_D_ (colored lines matched to islet color) for each islet was modeled over the course of low and high glucose perfusion (black dashed line), showing variable islet responses depending on location within the chamber and proximity to perfused blood vessels.

### Incorporating EC and stromal cells into islets enhances islet vasculature

Given the importance of vessel proximity to islets demonstrated by our COMSOL model, we tested whether increasing intra-islet vasculature in the islet-VMO might improve glucose responsiveness of embedded islets. We therefore dissociated native islets and then reconstituted them, either with islet cells alone (“reaggregated islets”) or with ECs and islet-derived stromal cells (“pseudo-islets”) (Fig. 6A). Upon reconstitution, reaggregated islets have poorly defined boundaries (Fig. 6B-C), whereas the inclusion of EC and stromal cells produced a more clearly defined boundary, indicating a more compact islet structure (Fig. 6D). Static GSIS assays (Fig. 6E) revealed that reaggregated islets have higher basal and stimulated secretion, although the stimulation index was reduced relative to native islets (2.0 vs. 4.7-fold, respectively). Compared to reaggregated or intact islets, basal insulin secretion and the stimulation index (2.7) were intermediate in pseudo-islets.

**Figure 6.**
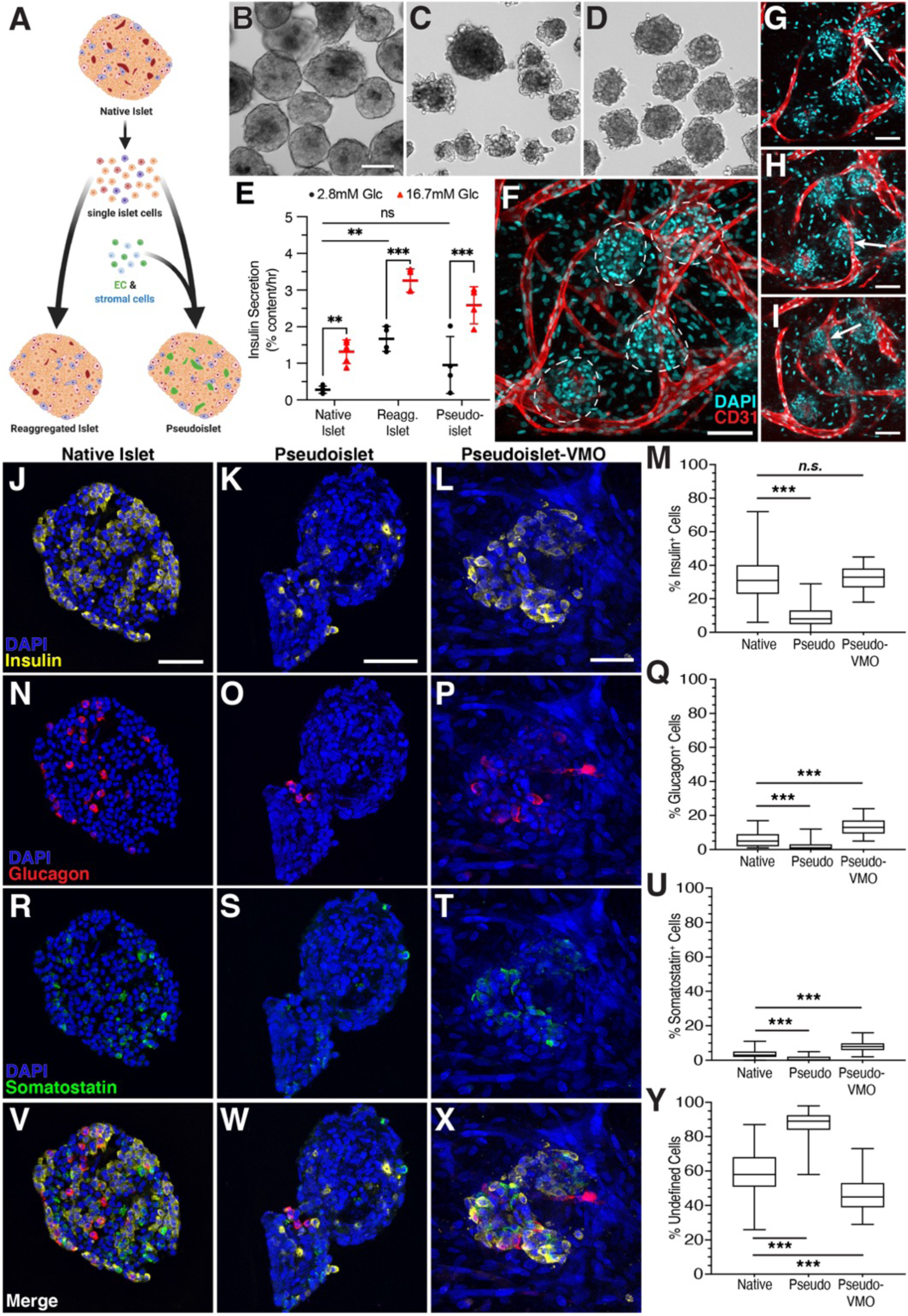
Intra-islet vasculature is enhanced in pseudo-islets embedded within the islet-VMO. (A) To generate reconstituted islets, native islets were dissociated and reconstituted either as islet cells alone (reaggregated islets) or together with EC and stromal cells (pseudo-islets). Image created with BioRender.com. (B-D) Islet morphology was compared by phase contrast microscopy two days post-reconstitution between (B) native islets, (C) reaggregated islets, and (D) pseudo-islets. Scale bar, 100μm. (E) Static glucose-stimulated insulin secretion (GSIS) was measured for each islet population under unstimulated (2.8mM glucose) and stimulated (16.7mM) conditions (two-way ANOVA, significance: n.s., not significant; **p<0.01; ***p<0.001). Error bars represent standard deviation. (F) Pseudo-islets were loaded together with EC and stromal cells and vessel networks allowed to form. ECs (CD31^+^, red) form an interconnected network that surrounds and penetrates the pseudo-islets (DAPI, cyan, dashed outline). Scale bar, 100μm. (G-I) Optical slices show vessels penetrating pseudo-islets (white arrows). Scale bar, 100μm. Immunofluorescent staining and quantification of native islets, pseudo-islets after reconstitution, and pseudo-islets maintained in the islet VMO for one week (pseudoislet-VMO) for (J-M) insulin^+^, (N-Q) glucagon^+^, and (R-U) somatostatin^+^ cells. Box and whisker plots represent median, 25^th^, and 75^th^ percentiles (box) and min and max values (whiskers) for each data set. Scale bar, 50μm. Student’s t-test, n.s., not significant; ***p<0.0001). (V-X) Merged images of staining in all islet types and (Y) quantitation of undefined (non-endocrine) islet cells.

To determine whether incorporation of EC and stromal cells into the pseudo-islets facilitates improved intra-islet vasculature in the islet-VMO platform, pseudo-islets were co-loaded along with additional EC and stromal cells into the platform two days post-reconstitution. We found blood vessel formation not only next to pseudo-islets, but also inside the islets (Fig. 6F-I). In the VMO platform, the pseudo-islets showed a similar number of insulin^+^ cells as native islets, whereas these cells were considerably diminished in number in the pseudo-islets cultured in suspension (Fig. 6J-M). Interestingly, while the number of glucagon^+^ and somatostatin^+^ cells was also decreased in suspension pseudo-islets compared to native islets, both populations were significantly enriched in pseudo-islets in the VMO (Fig. 6N-Q, 6R-U). Overall, pseudo-islets in suspension had a greatly increased number of undefined (non-endocrine) cells compared to the other groups (Fig. 6V-Y).

Interestingly, insulin release was not detected from pseudo-islets within the islet-VMO (data not shown). Previous work demonstrating that both *α* and *δ* cells can regulate β cell insulin release (36) raises the possibility that imbalance of these endocrine cell types underlies defective insulin secretion from the pseudo-islet-VMO, which is being further investigated.

### Perfusion of islet-activated immune cells to model islet-immune cell interactions

To assess the utility of our platform for investigating immune cell interactions with islets, we developed a non-autologous system for proof-of-concept studies. We cultured non-MHC matched PBMC with freshly isolated islets either alone or in the presence of blocking antibodies to MHC class I and II (Fig.7A). IFN-*γ* was added to increase MHC molecule expression and IL-2 was added to drive proliferation of any T cells that became activated. As expected, a sub-population of the T cells in the PBMC population had T cell receptors that cross-reacted with the (foreign) islet MHC, leading to allogeneic activation (indicated by the presence of grape-like clusters of cells), which was blocked by the anti-MHC antibodies (Fig. 7B,C). Proliferation was confirmed by cell counting (Fig. 7D).

**Figure 7.**
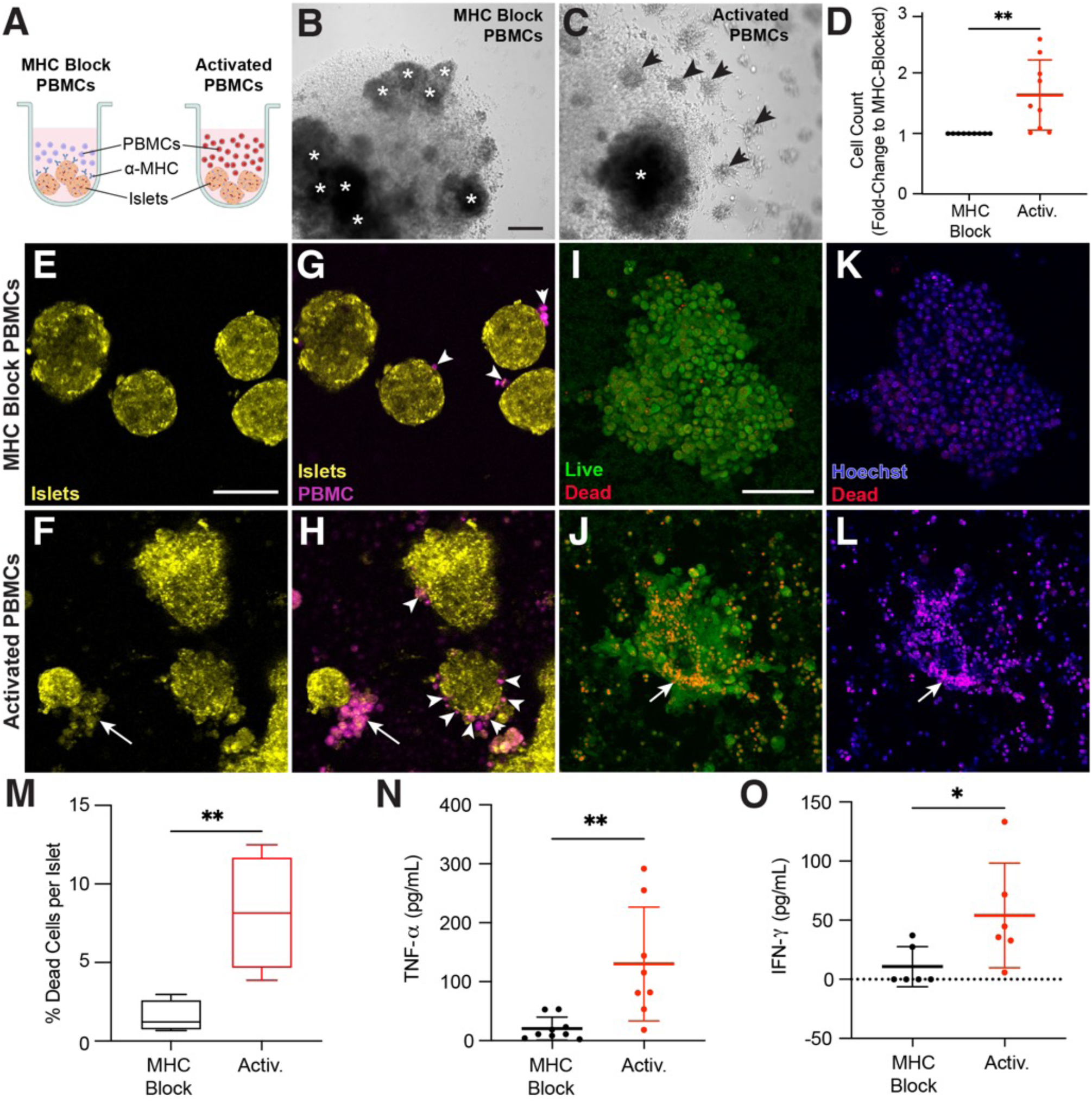
PBMCs are activated against donor islets. (A) MHC-blocked and activated PBMCs were generated by incubating PBMCs with donor islets in the presence of IL-2 and IFN-g (activation cocktail) in round-bottom 96-well plates. MHC-blocked conditions contained antibodies against both MHC Class I and II. Schematic created with BioRender.com. After 5 days under these conditions, PBMCs cultured in (B) the presence of, or (C) the absence of MHC-blocking antibodies demonstrates clonal expansion (activation) only in the absence of MHC-blocking antibodies. Islets are annotated by asterisks (*) and grape-like clusters annotated by arrowheads. Scale bar, 200mm. (D). Quantification of activated PBMCs (red) as a fold-change relative to MHC-blocked PBMCs (black) after five days of activation. The MHC-blocked and activated PBMCs were isolated and incubated with islets from the same donor for 48 hours. (E-H) Islets (yellow) and PBMCs (magenta) were stained with different CellTracker dyes and imaged for interactions between the two after 48 hours incubation in a chamber slide. Islets exposed to activated but not MHC-blocked PBMCs show morphological damage (E, F). (G) MHC-blocked PBMCs show minimal affinity for islets (white arrowheads) whereas, (H) activated PBMCs show increased adhesion to islets and, in some cases, destruction of islet tissue (white arrows). (I-L) Islets from the same conditions were stained with live/dead stain. (I, J) Morphological damage is evident in islets exposed to activated but not MHC-blocked PBMCs (arrow indicates islet damage). (K, L) Increased dead cells (red) in islets exposed to activated but not MHC-blocked PBMCs. (M) Quantification of dead cell labeling (red) versus Hoechst (nuclei, blue) staining in islets exposed to MHC-blocked (black) or activated (red) PBMCs (Student’s t-test, **p<0.01) (n=30 islets across 4 donors). Supernatant was collected from these co-incubations and measured for secreted TNF-a (N) and IFN-g (O). Statistical significance determined by Student’s t-test (significance: *p<0.05, **p<0.01).

To determine the killing capacity of the activated cell population, MHC-blocked and activated cells were stained with CellTracker Green then incubated with CellTracker Red-stained whole islets from the same donor used for the initial stimulation. Live cell imaging after two days showed morphological damage to islets exposed to activated immune cells, but not those incubated with MHC-blocked cells (Fig. 7E-F). Moreover, MHC-blocked PBMC infrequently interacted with islets, whereas activated cells regularly surrounded islets (Fig. 7G-H). Examination of islets with live/dead staining shows that islets exposed to activated, but not MHC-blocked, cells experience morphological damage (Fig. 7I-J). Islets exposed to activated PBMC also have a higher percentage of dead cells than those exposed to MHC-blocked cells (Fig. 7K-M). ELISA analysis of the supernatant from these assays also shows increased secretion of the inflammatory cytokines TNF-*α* and IFN-*γ* from activated cells interacting with the islets (Fig. 7N-O).

Having found that islet-reactive lymphocytes can be generated in response to allogeneic islets, we next sought to determine whether these cells can traffic to islets within the islet-VMO. MHC-blocked or activated PBMC were added to reservoir V_A_ and perfused through platforms containing islets from the same donor (Fig. 8A). After 48 hours of perfusion we noted robust extravasation of activated PBMC, whereas we saw very few MHC-blocked PBMC entering the tissue (Fig. 8B-C, S3A-B). Many of the extravasated cells migrated towards the islets, with some also visible within the islet structure itself (Fig. 8D). Indeed, activated PBMC demonstrated significantly higher rates of adhesion and extravasation compared to MHC-blocked PBMC (Fig. 8E). To determine if activated PBMC preferentially traffic to (or are retained in) islets, the number of immune cells within 100μm of an islet (Adjacent - Adj) was quantified and compared to control (Fig. 8F). MHC-blocked immune cells showed no preference for islets compared to background regions (Fig. 8F). In sharp contrast, immune cells that had been pre-activated against islet MHC showed a strong and significant bias toward islet localization (Fig. 8F). To determine if activated PBMCs also *invade* islets, the number of immune cells within an islet was quantified and compared to cells in background areas and to MHC-blocked cells in islets. Again, compared to the control cells, the activated cells were strongly biased toward islet invasion (Fig. 8G). To confirm that it is indeed T cells invading the islets, we stained tissues for CD3 and found numerous CD3^+^ cells both in and around islets in tissues perfused with activated PBMC, which contrasts with the very few CD3^+^ cells present in tissues perfused with control (MHC-blocked) PBMC (Fig. 8H-K). Thus, allogeneic T cells with specificity for islets can be introduced through the vasculature of the islet-VMO platform where they can extravasate and migrate into islets.

**Figure 8.**
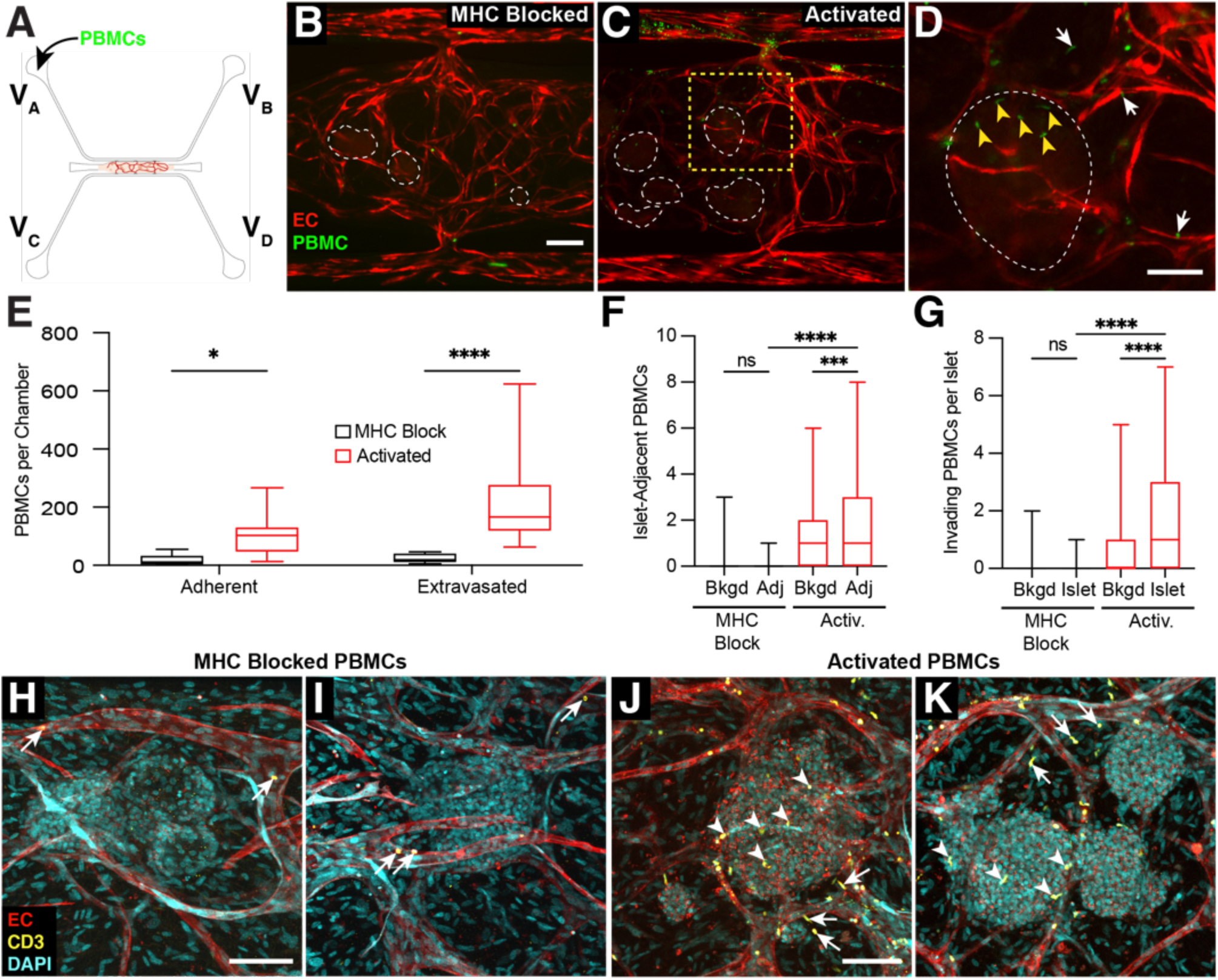
Immune cell interactions with islets in the islet-VMO. (A) CellTracker^+^ MHC-blocked or activated PBMCs were added to reservoir V_A_ and allowed to perfuse though islet-VMOs containing islets from the same donor for up to 48 hours. (B) MHC-blocked PBMCs (green) demonstrate minimal adhesion or extravasation from vessels (fluorophore-transduced ECs, red) as compared to (C) activated PBMCs (islets, dashed outline). Scale bar = 200mm. (D) Inset from (C) (yellow dashed box) shows increased migration of activated PBMCs out of the vasculature towards the islet (white arrows) with multiple PBMCs co-localizing within the islet itself (yellow arrowheads). (E) Quantification shows increased adhesion and extravasation of activated but not MHC-blocked PBMCS across multiple islet donors (two-way ANOVA, *p<0.05, ****p<0.0001) (n=14 Islet-VMOs across 4 islet donors). (F) The number of PBMCs within 100mm of an islet was quantified (Adj) and compared to both background, non-islet regions (Bkgd) and to MHC-blocked PBMCs (one-way ANOVA, ***p<0.001, ****p<0.0001 by) (n>65 islets across 4 islet donors). The number of PBMCs present within islets (islet) was quantified and compared to PBMC counts in background, non-islet regions (Bkgd) and to MHC-blocked PBMCs (one-way ANOVA, ***p<0.001, ****p<0.0001 by) (n>65 islets across 4 islet donors). (H, I) Confocal imaging of fixed and immunofluorescently-stained islet-VMOs perfused with MHC-blocked PBMCs shows minimal adhesion (arrows) of CD3^+^ T cells (green) to the vessels (CD31^+^, red; DAPI, cyan). (J, K) Immunofluorescent staining of islet-VMOs perfused with activated PBMCs shows increased extravasation (white arrows) and islet (DAPI, cyan) invasion (yellow arrowheads) of CD3^+^ T cells. Scale bars, 100mm.

## DISCUSSION

Here we describe a novel, biomimetic approach to recapitulate aspects of the native human islet environment while maintaining islet function *ex vivo*. Specifically, the islet-VMO uniquely incorporates (a) a 3D microenvironment comprised of ECM and human stromal cells mimetic of the native pancreas, (b) perfusable human blood vessels that transport glucose and insulin to and from islets, and (c) the capability to deliver immune cells via this vasculature. Importantly, the islets maintain glucose responsiveness for at least a week in the platform. Others have explored some of these strategies for modeling the native human islet environment, including recapitulating the surrounding ECM (37; 38), providing physiologically-relevant interstitial flow (39–41), and co-culturing with ECs (42). The islet-VMO, however, incorporates all of these features with the addition of a perfusable human vasculature, resulting in a uniquely powerful tool capable of reproducing physiologically-relevant islet functions and immune interactions.

The kinetics of insulin secretion we see with the islet-VMO are quite different from that of standard perifusion assays, where human islets exhibit biphasic GSIS characterized by a brief, high-amplitude first phase, following by a lower, sustained second phase (34). In contrast, in the islet-VMO we see a more gradual slope of first-phase insulin release and the absence of a distinct second phase. Previous work has also shown the lack of a second-phase response when islets are encapsulated in a hydrogel (43), consistent with our findings. Our COMSOL modeling also suggests that the islet response is determined both by location within the chamber and proximity to perfused vessels. Critically, the insulin secretion dynamics demonstrated in the islet-VMO better represent physiological insulin secretion *in vivo* (Fig. 3), which is also characterized by a gradual accumulation of circulating insulin without an apparent second phase of insulin release (35). Remaining differences in insulin secretion dynamics between the islet-VMO and the *in vivo* setting highlight opportunities for further improvements in modeling physiological insulin secretion, such as incorporating islet innervation and co-stimulating with incretin hormones.

Diabetes is a complex disease characterized by both molecular and immunological triggers that drive disease pathogenesis. Interaction between immune cells and islets is important in both forms of diabetes, wherein T1D is driven by autoimmune reactivity of T cells against β cells (44) while T2D is characterized by β cell inflammation and activation of tissue resident and circulating macrophages (3; 6; 45). In this study, we have demonstrated the feasibility of using the islet-VMO to understand these immune cell interactions. The model recapitulates physiological immune cell delivery through vessels, trafficking (extravasation), and islet invasion. Importantly, while these processes can be modeled in mice, the islet-VMO exclusively utilizes human cells in the tissue and immune environment, thereby improving the relevance of discoveries for human patients.

A fully autologous human system – that captures patient-specific genetic backgrounds – is a highly-desired tool in the field of diabetes research. We and others have achieved several milestones towards generating such a platform. First, recent work demonstrates the feasibility of deriving all of the relevant cell populations from iPSCs, including islet cells (46; 47), ECs (48), and stromal cells that can be paired with patient-matched immune cells. Second, β cells can be generated from iPSCs derived from T1D patients (49), enabling incorporation of β cells with susceptible genetic backgrounds into the platform. Third, in this work we have demonstrated an approach to generate pseudo-islets from dissociated cell populations and then incorporate these into a vascularized islet-VMO. This capability will be key for incorporating iPSC-derived cells into the VMO system. Achieving glucose responsiveness from pseudo-islets in the VMO, which is a goal of our ongoing work, would further expand the capability to reconstitute the native islet environment. Lastly, a similar but simpler 2D approach using iPSC-derived β cells and patient-matched PBMC from both T1D patients and healthy volunteers has demonstrated the feasibility of using these cells to model relevant cell-cell interactions (50). We anticipate that the addition of perfused vasculature in the islet-VMO will further build on this model to provide more physiologically-relevant interpretations of diabetes pathogenesis in the laboratory.

## ACKNOWLEDGEMENTS

***Financial Support.*** This work was supported by funding from the National Institutes of Health (#UC4DK104202-01 & #UH3DK122639-03 to C.C.W.H. and M.S.), Juvenile Diabetes Research Foundation postdoctoral fellowship 3-PDF-2014-193-A-N (M.W.), 3-PDF-2017-386-A-N (K-V.N-N.), 3-PDF-2020-932-A-N (Y.J.), and John G. Davies Endowed Fellowship in Pancreatic Research S1105-1002847-AWD (M.W.). Human cadaveric islets and relevant donor information were supplied by the Integrated Islet Distribution Program (IIDP, NIH Grant #2UC4DK098085).

## Author Contributions

R.H.F.B. conceptualization, methodology, investigation, data curation, formal analysis, validation, visualization, supervision, writing – original draft. B.T.O. methodology, investigation, data curation, formal analysis, software, validation, visualization, writing – review & editing. B.S. methodology, investigation, data curation, formal analysis, software, validation, visualization, writing – review & editing. B.Q.P. investigation, formal analysis, writing – review & editing. D.J.J. investigation, writing – review & editing. M.S.H. investigation, writing – review & editing. V.S.S. software, writing – review & editing. M.W. methodology, investigation, writing – review & editing. K-V.N-N. methodology, investigation, writing – review & editing. Y.J. methodology, investigation, writing – review & editing. R.G. methodology, writing – review & editing. K.L.C. funding acquisition, methodology, writing – review & editing. L.T. methodology, writing – review & editing. S.C.G funding acquisition, methodology, data curation, formal analysis, writing – review & editing. M.S. conceptualization, funding acquisition, methodology, supervision, project administration, resources, writing – review & editing. C.C.W.H. conceptualization, funding acquisition, methodology, supervision, project administration, resources, writing – review & editing.

