## Supplemental Materials for "A vascularized 3D model of the human pancreatic islet for *ex vivo* study of immune cell-islet interaction"

**SUPPLEMENTAL FIGURE LEGENDS**

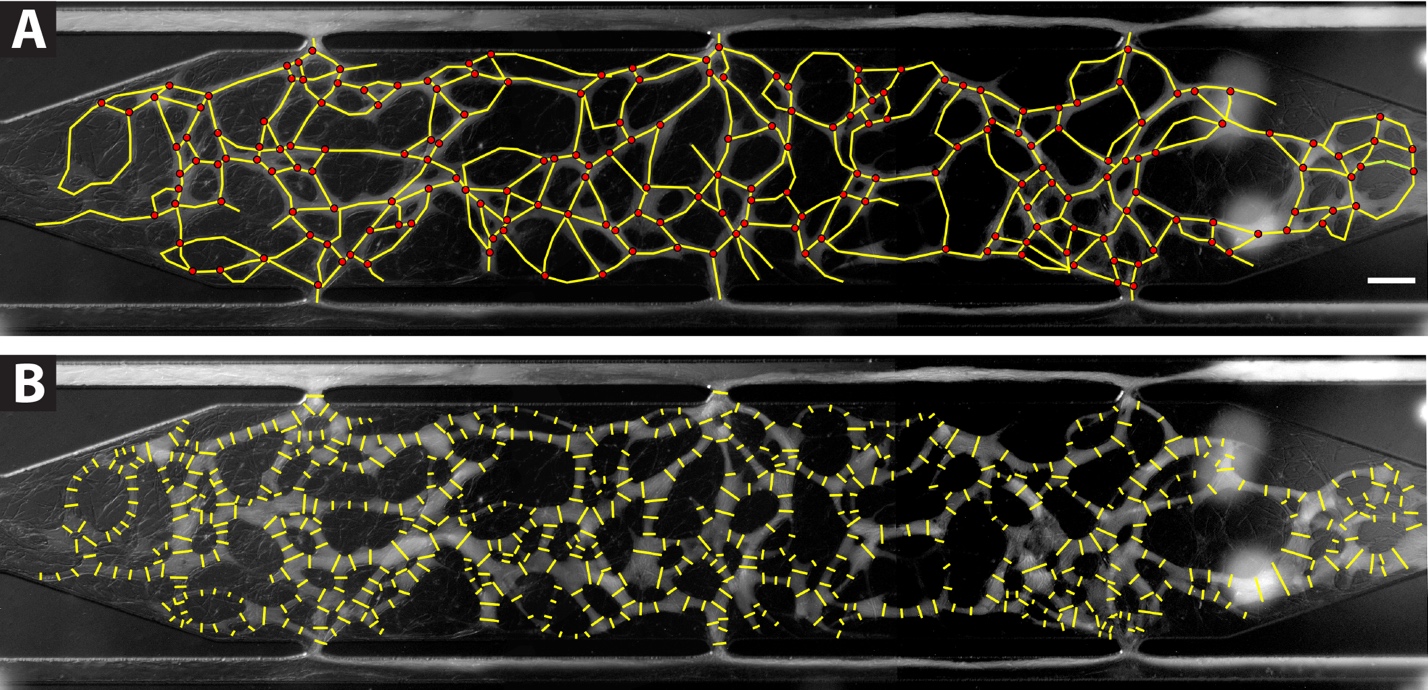
**Figure S1. Measuring vessel network parameters in the islet-VMO.** (**A**) Vessel length was measured using FIJI imaging software by measuring the length of segmented lines (yellow) traced at the midpoint of all vessels. Branching was quantified by counting the intersection points of these lines (red circles) throughout the cell chamber. Scale bar, 200μm. (**B**) To measure vessel diameter, the cross-section of all traced vessels was measured at 50-100μm intervals along the length of each vessel.

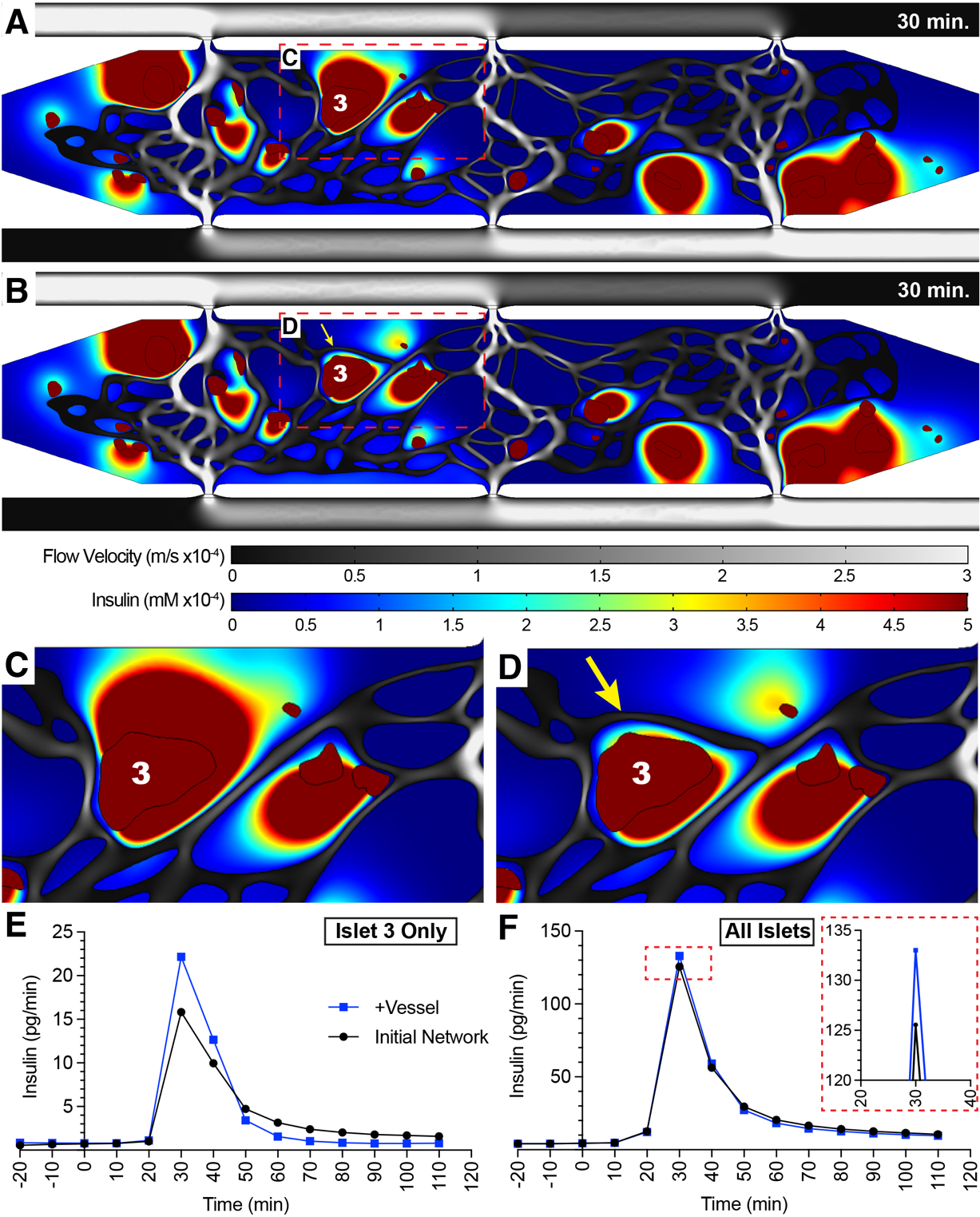

**Figure S2. Insulin secretion of individual islets is dependent on proximity to perfused blood vessels.** Insulin secretion was first modeled with the skeletonized vessel network from Fig. 3, showing insulin accumulation around islet #3. A snapshot of flow velocity (m/s, gray scale) and insulin secretion (mM, color scale) are plotted 30 minutes post-high glucose perfusion. (B) Addition of a new connecting blood vessel (yellow arrow) reduces the size of this insulin pool, demonstrating the importance of vessel location for trafficking secreted insulin. (C, D) Insets of before (A) and after vessel addition (B) demonstrates the reduced insulin accumulation in the presence of the new vessel. (E) Quantification of the insulin contribution from islet #3 alone before vessel addition (black line) and after vessel addition (blue line). (F) Quantification of total insulin production from the entire chamber before vessel addition (black line) and after vessel addition (blue line).

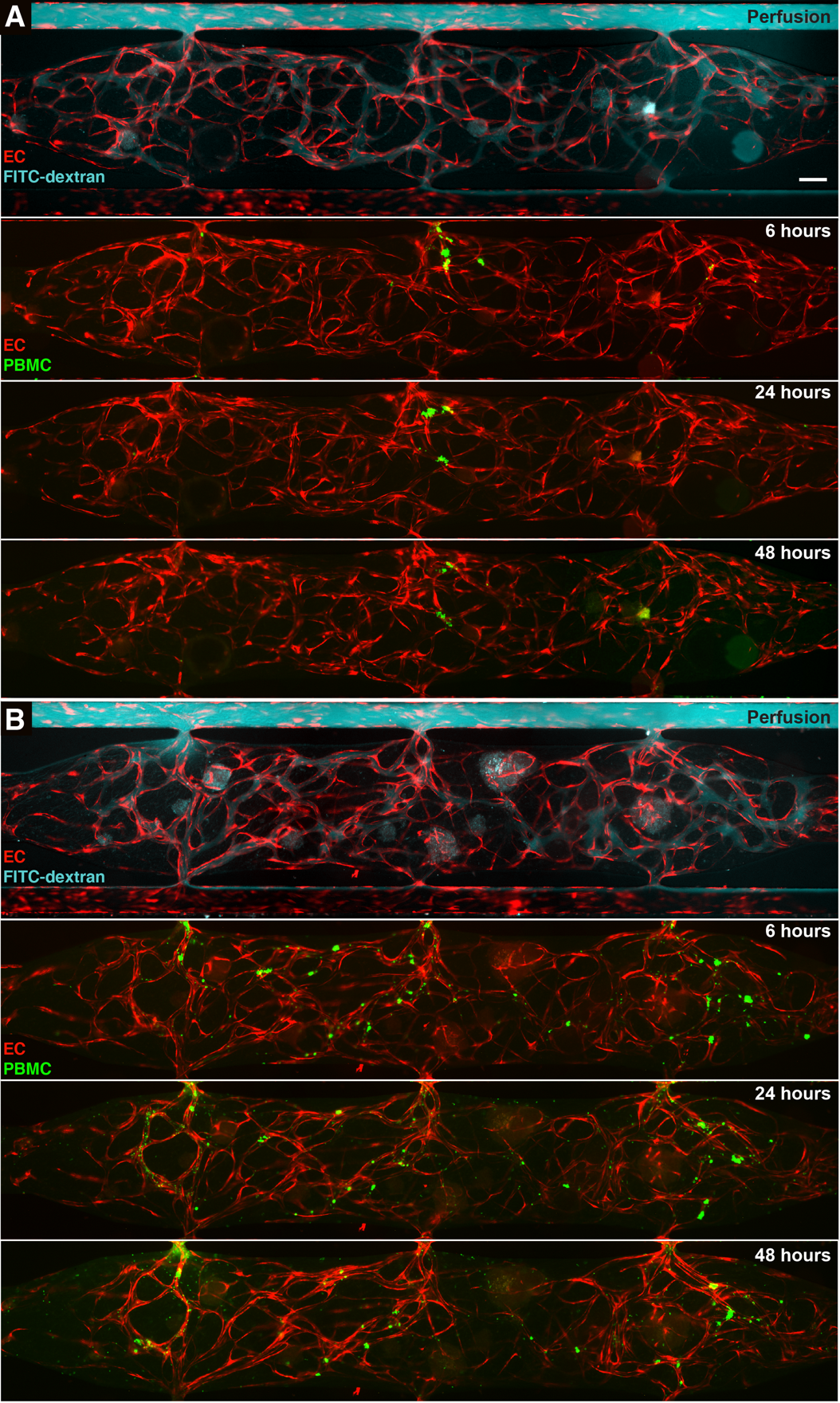

**Figure S3. Time course of PBMC perfusion through the islet-VMO.** Adhesion and extravasation of (**A**) MHC-blocked PBMCs (green) from blood vessels (fluorophore-transduced ECs, red) is reduced at multiple timepoints as compared to (**B**) activated PBMCs at the same time points. Serial images within each panel demonstrate dextran perfusion (prior to PBMC perfusion), and 6, 24, and 48 hours post-PBMC perfusion. Scale bar, 200μm.

**SUPPLEMENTAL TABLES**

**Table S1. Medium Formulations.**

| **Medium Name** | **Formulation** |
| --- | --- |
| Islet Medium | 10% FBS, 13mM glucose, 2mM L-glutamine, 1mM sodium pyruvate, 10mM HEPES, 0.25µg/mL amphotericin B, 100U/mL PenStrep, CMRL 1066 medium |
| Krebs-Ringers Buffer (KRB) | 130mM NaCl, 5mM KCl, 1.2mM CaCl_2_, 1.2mM MgCl_2_, 1.2mM KH_2_PO_4_, 20mM HEPES pH 7.4, 25mM NaHCO_3_, 0.1% BSA, 2.8mM glucose in water |
| M199(+) | M199 (-phenol red) medium, EGM-2 growth factors (Lonza Bioscience), 10ng/mL VEGF-165 (Shenandoah Biotechnologies, Warminster PA) |
| High-Glucose M199(+) | M199(+), 16.7mM glucose (final) |
| PBMC Medium | RPMI-1640 medium, 10% FBS |
| PBMC Activation Medium | RPMI-1640 medium, 10% FBS, 10ng/mL IL-2 (Biolegend), 20ng/mL IFN-γ (Biolegend) |

**Table S2. Antibodies for immunofluorescence (IF) staining, flow cytometry (FC), and blocking (B).**

| **Antibody** | **Species** | **Application** | **Supplier** | **Catalog #** | **Dilution** |
| --- | --- | --- | --- | --- | --- |
| CD31 | mouse | IF | Agilent Technologies Inc., Santa Clara CA | #M0823 | 1:600 (IF, section), 1:300 (IF, VMO) |
| CD3 | rabbit | IF | Abcam, Cambridge MA | #ab135372 | 1:300 (IF, VMO) |
| Glucagon | mouse | IF | Sigma-Aldrich, St. Louis MO | #G2654 | 1:5000 (IF, section) 1:1,000 (IF, VMO) |
| Insulin | guinea pig | IF | Agilent Technologies Inc., Santa Clara CA | #A056401 | 1:1,000 (IF, section) 1:500 (IF, VMO) |
| HLA-A, B, C | Mouse | B | BioLegend, San Diego CA | #311402 | 10μg/mL |
| HLA-DR | Mouse | B | BioLegend, San Diego CA | #307602 | 10μg/mL |
| Laminin 1+2 | rabbit | IF | Abcam, Cambridge MA | #ab7463 | 1:100 (VMO) |
| Somatostatin | goat | IF | Santa Cruz Biotechnology Inc., Dallas TX | #sc7819 | 1:500 (IF, section) 1:300 (IF, VMO) |

**Table S3. Constants for COMSOL Modeling.**

| **Parameter** | **Constant** | **Source** |
| --- | --- | --- |
| First-phase insulin release, Hill-slope* | 4 | (1; 2) |
| Second-phase insulin release, Hill-slope* | 0.1 | (1; 2) |
| Glucose consumption, Hill-slope | 3 | (1; 2) |
| First-phase insulin response | 3.43 x 10^-4^ mol/m^3^/s | (1; 2) |
| Second-phase insulin response | 5.15 x 10^-6^ mol/m^3^/s | (1; 2) |
| Oxygen diffusion coefficient in fibrin | 1.7 x 10^-9^ m^2^/s | (3) |
| Insulin diffusion in fibrin | 8 x 10^-12^ m^2^/s | Fluorescent insulin, fluorescent recovery after photobleaching (FRAP) (data not shown) |
| Glucose diffusion in fibrin | 3 x 10^-10^ m^2^/s | (4-8) |
| Fibrin permeability | 1.5 x 10^-13^ m^2^ | (9) |
| Endothelial cell hydraulic conductivity | 3.5 x 10^-9^ cm/s*Pa | (10) |
| **Alterations were made to Hill-slopes and insulin production rates in response to the insulin output kinetics of the islet-VMO.* | | |
